# Genome wide association study reveals novel candidate genes associated with productivity and disease resistance to *Moniliophthora* spp. in cacao (*Theobroma cacao* L.)

**DOI:** 10.1101/820944

**Authors:** Jaime A. Osorio-Guarín, Jhon A. Berdugo-Cely, Roberto A. Coronado-Silva, Eliana Baez, Yeirme Jaimes, Roxana Yockteng

## Abstract

Cacao (*Theobroma cacao* L.), the source of chocolate, is one the most important commodity products for farmers to improve their economic benefits. However, diseases such as frosty pod rot (FPRD) caused by *Moniliophthora roreri* and witches’ broom (WBD) caused by *Moniliophthora perniciosa*, limits the increase in yields. Molecular tools can help to develop more rapidly cacao varieties with disease resistance. In the present study, we sequenced by genotyping-by-sequencing (GBS), 229 cacao accessions to examine their genetic diversity and population structure. From those accessions, 102 have been evaluated for disease resistance and productivity to conduct a genome-wide association study (GWAS) based on 9,003 and 8,131 SNPs recovered by mapping against to the annotated cacao genomes (Criollo and Matina). Three promissory accessions for productivity and 10 accessions showing good tolerance to the evaluated diseases were found in the phenotypic evaluation. The work presented herein provides the first association mapping study in cacao using SNP markers based on GBS data. The GWAS identified two genes associated to productivity and seven to disease resistance. The results enriched the knowledge of the genetic regions associated to important traits in cacao that can have significant implications for conservation and breeding strategies such as marker-assisted selection (MAS).

## INTRODUCTION

The cacao species (*Theobroma cacao* L) is a perennial crop and represents one of the oldest agroforestry systems in tropical America (Bergmann 1969). The geographical origin of cacao is Tropical South America (Motamayor *et al*. 2002), where several wild populations can be found in the Amazon, Orinoco, and Guyana regions (Motamayor *et al*. 2008; Zhang *et al*. 2012). The cacao beans are the primary source of the multibillion-dollar industry that produces chocolate and the primary income for about 6 million smallholder farmers globally (World Cocoa Foundation 2014).

In Colombia, cacao cultivation occupies an area of 177 thousand hectares with approximately 38,000 farmer families and generates more than 155,000 jobs (MADR 2018). The country is classified as the tenth producer worldwide with a yield close to 400 kg/ha and a production of 54,000 metric tons of beans (Ríos *et al*. 2017). The top export destinations of Colombian production are Mexico, Italy, Spain, and Netherlands (Castro and Vignati 2018). Predictions of the cocoa market estimate that the demand will increase by 30% in 2020, forcing to cacao producer countries to increase their production (Fountain and Hütz-Adams 2018). The most important limiting factors for cacao production worldwide are diseases caused by fungal and oomycete pathogens, up to 40% of the crop production is estimated to be lost annually due to diseases (Bowers *et al*. 2001; Álvarez *et al*. 2014).

Among the main diseases that attack cacao are frosty pod rot (FPRD) commonly known as monilia and witches’ broom (WBD) caused by the oomycetes *Moniliophthora roreri* and *Moniliophthora perniciosa,* respectively (Bailey and Meinhardt 2016). Several disease control methods have been attempted to reduce the incidence of FPRD and WBD from which the cultural practices are the most used methods. Another control method is the application of fungicides that affect the physiological processes of the causal agent. The biological control is also used and is based on the application of microorganisms or their exudates or molecules to generate antibiosis and stop the growth of the pathogen (Bateman *et al*. 2005; Loguercio *et al*. 2009; Acebo-Guerrero *et al*. 2012). The breeding strategy is the development of varieties with disease resistance that could reduce the dependence on chemical substances and the high labor costs of cultural control (Jaimes and Aranzazu 2010).

The development of resistant or high-yielding varieties could be accelerated by using marker-assisted selection (MAS), which refers to the selection of individuals based on molecular markers associated to resistance/defense quantitative trait loci (QTL) (Collard and Mackill 2008). During recent decades, numerous QTL for productivity (Crouzillat *et al*. 2000; Clement *et al*. 2003) and resistance to FPRD or WBD (Brown *et al*. 2005; Faleiro *et al*. 2006; Queiroz *et al*. 2008; Motilal *et al*. 2016) have been identified using bi-parental populations. Based on the SNP array data, Royaert *et al*. (2016) described genomic regions involved in WBD resistance and McElroy *et al*. (2018) and Romero-Navarro *et al*. (2017) identified several markers related to productivity and disease resistance to *Moniliophthora* spp.

The constant advances in next-generation sequencing technologies have allowed the possibility to generate thousands of single nucleotide polymorphisms (SNPs) by reducing genome complexity through methylation-sensitive restriction enzymes, technique known as genotyping-by-sequencing (GBS) (Elshire *et al*. 2011). Recently, Osorio-Guarín *et al*. (2018) optimized a GBS protocol for cacao to eliminate false calls of SNPs, which can also cause problems in the mapping of candidate genes. Also, the release of the cacao genome sequences has provided the information for rapid identification, functional, and structural characterization of many gene (Argout *et al*. 2010; Motamayor *et al*. 2013).

Our study is the first to use GWAS using GBS data in order to explore and identify new SNP variants distributed across the genome and associate them with important agronomic traits. The aims of the present work were: 1) assess the genetic diversity and population structure; 2) assess the phenotypic characterization for productivity and disease resistance to FPRD and WBD; 3) identify marker-trait associations, and finally 4) identify genomic regions that have undergone selection on the evaluated traits.

## MATERIALS AND METHODS

### Plant material

A total of 229 diverse cacao accessions from the national germplasm bank located in the research center La Suiza (7°22’12’’N - 73°11’39’’W) of the Colombian Corporation for Agricultural Research (Agrosavia) were used (Table S1). The collection was planted in 1998 under an agroforestry system (Arguello *et al*. 1999). Two to six clones per accession were planted in lanes with a distance of 2.5 m. The species trees *Cordia gerascanthus L.* and *C. alliodora* L. distributed in lanes every 12 m provides the shade. All plants are grown under the same standard agronomic practices.

### DNA isolation and sequencing analysis

Genomic DNA of the 229 accessions were extracted from young leaf tissue using the DNeasy Plant Mini Kit (QIAGEN, Germany). The DNA concentration and quality were estimated using Qubit Fluorometer v2.0 (Life Technologies, Thermo Fisher Scientific Inc) and by electrophoresis on 1% agarose. Genomic DNA was digested with 2.0 units of the *Bsa*XI enzyme ((N)_9_AC(N)_5_CTCC(N)_10_) (New England Biolabs) and 2.0 units of the *Csp*CI enzyme ((N)_10-11CAA_(N)_5_GTGG(N)_12-13_) (New England Biolabs). The library preparation was carried out according to the protocol proposed by Osorio-Guarín *et al*. (2018). The barcoded samples were pooled and sequenced on an Illumina HiScan SQ instrument in the Molecular Genetics Laboratory at the research center Tibaitatá (4°41’45”N - 74°12’12”W).

### SNP discovery

Raw sequence reads were checked for quality with FastQC (Andrews *et al*. 2012) and both ends were trimmed with Trim Galore v0.5.0 (Krueger 2018). Reads presenting a phred quality score below 25 and a sequence length below 60 bp were removed. Samples were aligned to the two reference genomes of *T. cacao*, Criollo B97-61/B2 v2 (Argout *et al*. 2010) and Matina 1-6 v1.1 (Motamayor *et al*. 2013), using the software BWA v0.7.17 (Li and Durbin 2009). SNPs were discovered and called using Picard Tools v2.18.9 (Institut|e Broad 2019) and GATK software v3.8.0 (McKenna *et al*. 2010). Finally, the SNPs were filtered with VCFtools v4.2 (Danecek *et al*. 2011) according to the pipeline proposed by Osorio-Guarín *et al*. (2018).

### *In silico* digest

To identify all restriction cut site positions for the combination of *Bsa*XI and *Csp*CI restriction enzymes, we used the restrict package from the software Emboss v6.5.7.0 (Rice *et al*. 2000) with the two references genomes (Argout *et al*. 2010; Motamayor *et al*. 2013). The total number of fragments along the genomes were ordered per chromosome, then were summed using a size range of 200-700 bp and finally plotted with the software R (R development core team 2008). The number of fragments produced with the *in silico* predictions were compared to the sequenced resulting alignments using the package genomecov from BEDtools v2.27.0 (Quinlan and Hall 2010).

### Genetic diversity, linkage disequilibrium, and population structure analyses

The observed heterozygosity (Ho), expected heterozygosity (He), and the polymorphism information content (PIC) were performed with Cervus v3.0.7 (Kalinowski *et al*. 2007). A cluster analysis using the neighbor-joining (NJ) method with 1,000 permutations was performed using the ppoppr package in R Software (Kamvar *et al*. 2014) and visualized with the software FigTree v1.4.2 (Rambaut 2014).

To determine the mapping resolution for the GWAS, a linkage disequilibrium (LD) analysis was performed using the software VCFtools v4.2 (Danecek *et al*. 2011) with a sliding window of 500 bp. The LD decay was determined using a loess regression generated from the plotting of pairwise LD (*r^2^*) and physical distance (bp) with a threshold value of 0.2. For chromosomes that presented significant associations, LD heatmaps were constructed with a custom R script.

Finally, the population structure was examined using a bayesian model clustering analysis in the software Structure v2.3.4 (Pritchard *et al*. 2000) with the following parameters: number of populations (K) set from 1 to 15, repeated 10 times, a burn-in period of 50,000 iterations and 100,000 Markov Chain Monte Carlo (MCMC) repeats. The software Clumpp v1.1.1 (Jakobsson and Rosenberg 2007) was used to estimate the degree of congruence between independent runs. Visualization of the results was done with the package ggplot on R software. Accessions with a score higher than 0.80 were assigned to a pure group, while those with a value lower than 0.80 were assigned to the admixture group. The K optimum was evaluated by the Evanno method (Evanno *et al*. 2005) using the software Structure Harvester (Earl and VonHoldt 2011).

### Phenotypic data and statistical analyses

Agrosavia’s cacao germplasm is conserved in a region with natural presence of inoculum of different pathogens allowing to evaluate the resistance of accessions. The number of healthy and infected pods were counted weekly using two to six clones of 102 accessions during four harvests periods from September of 2016 to May of 2018 (Table S2). With the resulting data, we calculated the area under the disease progress curve (AUDPC) for all harvest periods using the formula proposed by Shaner and Finney (1977):

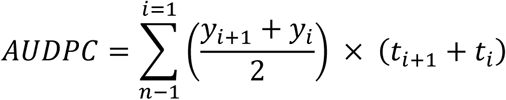

Where *yi* refers to the counting of the disease in the n^th^ observation; *ti* to the time in the n^th^ observation, and n to the total number of observations.

The following four variables were evaluated:

1. Healthy pods (productivity).
2. AUDPC of pods infected by FPRD.
3. AUDPC of flower cushion broom infected by WBD.
4. AUDPC of deformed branches infected by WBD.

The results were compared among the 102 accessions through an analysis of variance (ANOVA). Correlations among traits were calculated using Pearson’s correlation coefficient (*r*) at *p ≤ 0.05*. All statistical analysis were performed in R software (R development core team 2008).

We conducted a principal component analysis (PCA) with the prcomp package of the software R using the AUDPC values for each genotype. A cluster analysis using the Ward method and Euclidean distance was conducted using the two first components of the PCA.

### Association analysis

GWAS between the SNP markers and evaluated traits was performed for healthy pods (productivity) and AUDPC of the two diseases. The analysis was conducted using the genome association and prediction integrated tool (GAPIT) (Lipka *et al*. 2012), on the 102 accessions with phenotypic data. The mixed linear model (MLM) was used to discover the associations between polymorphisms and phenotypes and the risk of false association due to population stratification was minimized by incorporating population structure (Q) and kinship matrix (K) with the following formula:

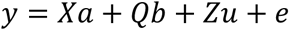

Where *y* is the vector for phenotypes; *a* is the vector of marker fixed effects, *b* is a vector of fixed effects, *u* is the vector of random effects (the kinship matrix), and *e* is the vector of residuals. X denotes the genotypes at the marker; Q is the Q-matrix, and Z is an identity matrix.

The Q-matrix was determined previously by Structure v1.3, and the kinship matrix was calculated in GAPIT with the Loiselle method. The quantile-quantile plots (Q-Q plots) were constructed to validate the appropriateness of the MLM. Manhattan plots were generated using the −log_10_(*_p_*) values for each trait.

### Identification of genomic regions under selection

The site frequency spectrum in the software SweeD (Pavlidis *et al*. 2013) and the pattern of LD between polymorphic sites was calculated with the software OmegaPlus (Kim and Nielsen 2004). The grid parameter was calculated for each chromosome in order to have a measure of the composite likelihood ratio (CLR) (SweeD) and ω (OmegaPlus) statistics every 10,000 bp. The minimum and the maximum size of the flanking region were fixed to 1,000 bp (min win option) and 100,000 bp (max win option). The common outliers were found using an R script that includes the annotation GFF file to identify the genes in the region under selection.

### Data availability

Table S1 contains the list of the accessions used in this study. The phenotypic data is provided in Table S2. Table S3 contains a summary of the statistics for the sequenced data per individual. The candidate genes under positive selection per chromosome are provided in Table S4. File S5 and S6 includes the vcf files of the SNPs discovered for Criollo and Matina, respectively. Figure S1 contains an *in silico* analysis of the restriction enzymes. Figure S2 is the correlation among traits of the whole population. Figure S3 contains the QQ-plots for each evaluated variable. Figure S4 presented the LD heatmaps for the chromosomes with significant associations. The R script for the detection of common outliers in selective sweeps is available at http://pop-gen.eu/wordpress/wp-content/uploads/2013/12/combined_analysis.zip. Supplemental material is available at Figshare: https://gsajournals.figshare.com/s/165b88f523482900eb79.

## RESULTS

### SNP Calling and *in silico* digestion

The sequencing of libraries generated by the GBS method resulted in 1,894 Gb raw reads and 1,555 Gb mapped reads. The observed read depth across samples ranged from 5.5 to 182.8 when mapped to the Criollo genome and from 7.1 to 179.6 when mapped to the Matina genome (Table S3). Genotyping of the 229 accessions without missing data, yielded a total of 9,003 SNPs compared to the Criollo genome, whereas 8,131 SNPs were found compared to the Matina genome.

To confirm that enzymes *Bsa*XI and *Csp*CI were suitable for the cacao genome, we previously performed an *in silico* digestion using the two reference genomes. The *in silico* analysis generated a higher number fragments with optimal sizes for sequencing in an Illumina system (200 to 700 bp) in Criollo genome (118,400) than in the Matina genome (42,849). This result indicates that the combination of the two enzyme patterns are more frequently found in Criollo. The number of base pairs was slightly lower in Criollo (8.5 million bp) than in Matina (9.1 million bp) because the produced fragments are longer for the latter (Figure S1).

The number of sequenced fragments produced per accession were lower than those predicted *in silico* for both genomes (Figure 1). When the reads were mapped against the two genomes, a higher number of fragments were recovered using the Criollo genome as reference.

**Figure 1.**
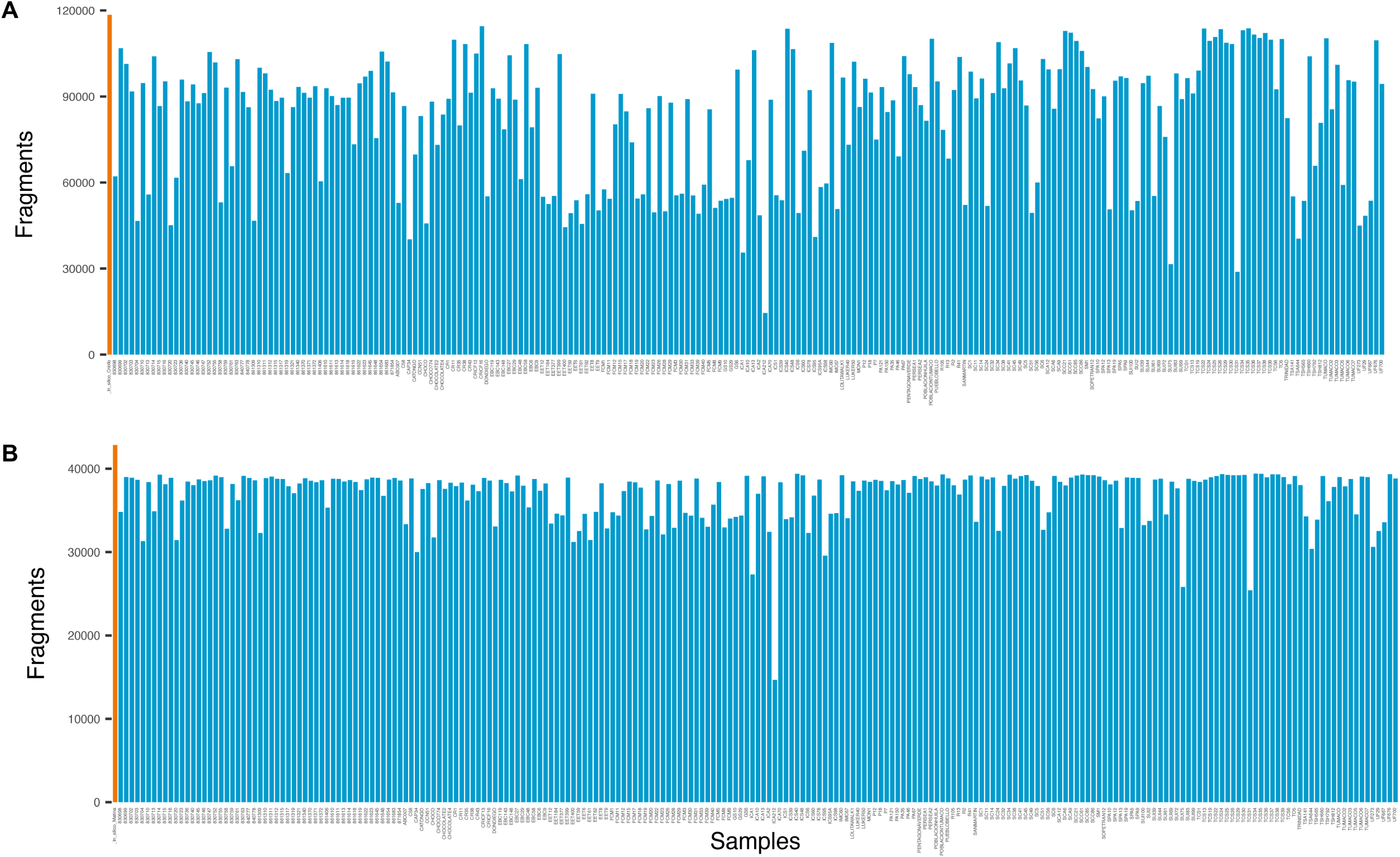
Number of sequenced reads with lengths between 200 and 700 bp of each accession mapped against **A.** Criollo genome and **B.** Matina genome. Orange bars correspond to the reads predicted with an *in silico* digestion of each reference genome.

### Genetic diversity

Values for He and Ho of the 229 cacao accessions were slightly higher with the dataset generated using the Criollo genome as reference than with the dataset generated using the Matina genome (Table 1). Medium to high levels of genetic diversity and an excess of heterozygosity were found in this collection. The PIC value showed that SNP markers were informative and was very similar for both datasets (Table 1).

**Table 1.** Genetic diversity statistics of the Agrosaviás cacao collection detected using the genomes Criollo and Matina of *T. cacao* as reference.

| Genome | Number of markers | Ho | He | PIC |
| --- | --- | --- | --- | --- |
| Criollo | 9,003 | 0.350 | 0.317 | 0.256 |
| Matina | 8,131 | 0.316 | 0.314 | 0.250 |

### Neighbor-joining, linkage disequilibrium and population structure analyses

The following analyses were carried out for the entire collection (229 accessions) using the dataset generated with the Criollo genome that contained the higher number of SNPs. The NJ analysis did not recover the 10 cacao genetic groups proposed by Motamayor *et al*. (2008); however, Agrosavia’s cacao germplasm formed subgroups than contained both modern landraces and accessions collected in wild regions (Figure 2).

**Figure 2.**
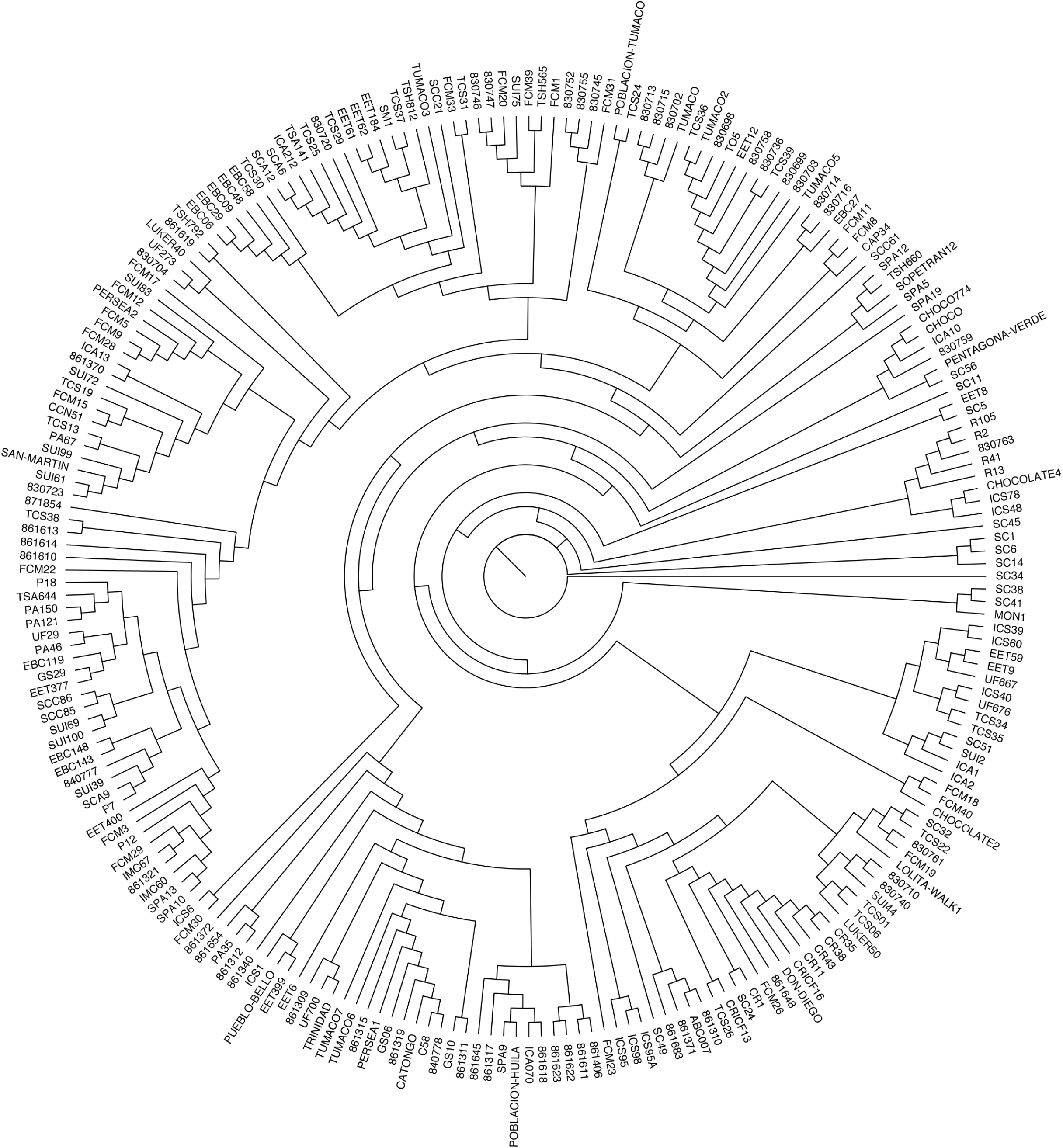
Neighbor-Joining tree based on Nei’s genetic distance of accessions of *T. cacao* of Agrosavia germplasm collection. Nei’s genetic distances were calculated using the 9,003 SNPs mapped against the Criollo genome.

The mean distance of LD decay was analyzed to characterize the mapping resolution for GWAS. The studied collection had an overall low LD and most of the *r^2^* values were below 0.25. In addition, LD declined rapidly with physical distance with an *r^2^* value of 0.125 within approximately 500 bp (Figure 3).

**Figure 3.**
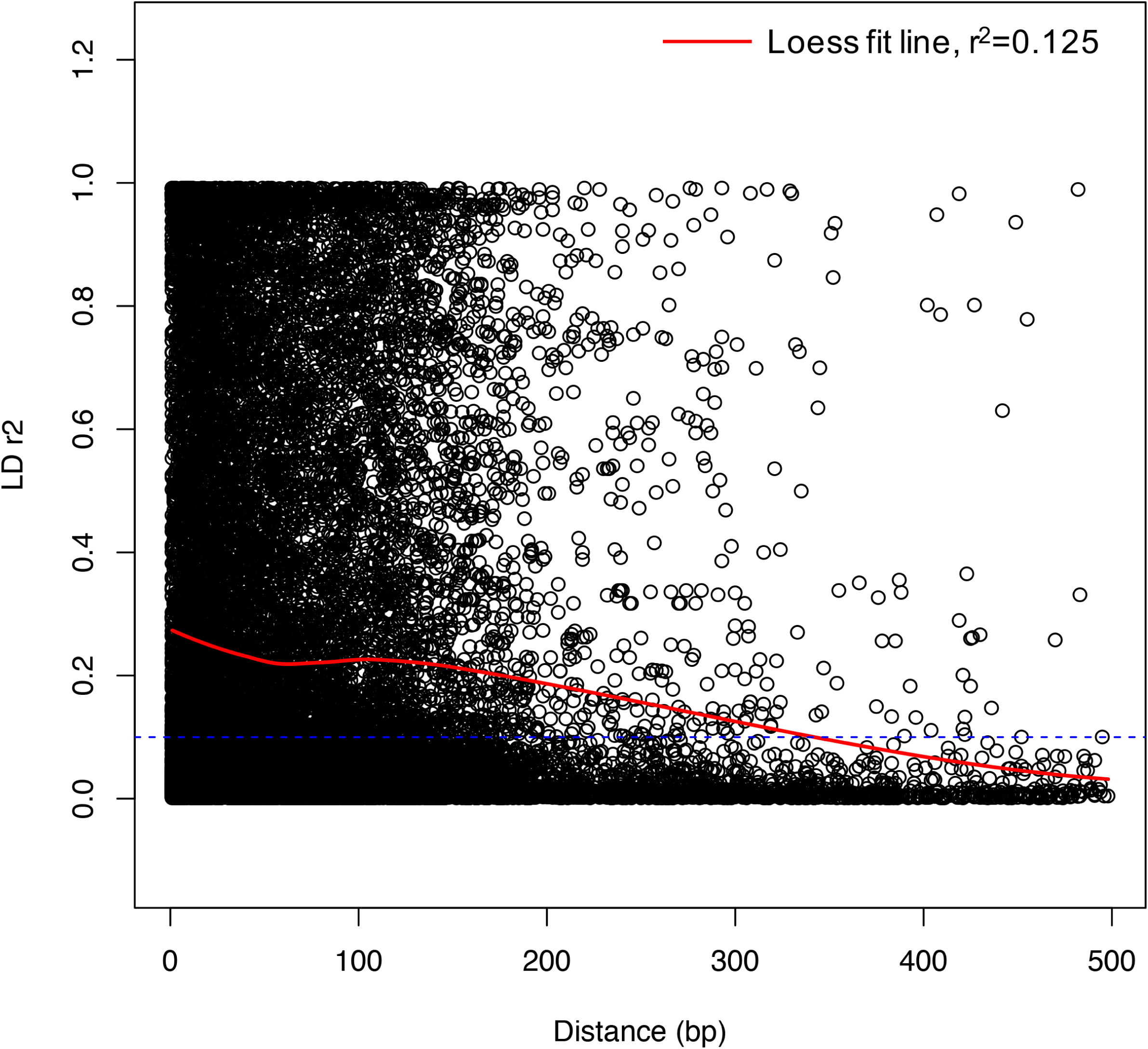
Analysis of linkage disequilibrium. LD Decay (*r*^2^) as a function of physical distance on all chromosomes. Only *r*^2^ values with *p ≤ 0.05* are shown.

Finally, the population structure showed that the best value for the number of sub-populations was 7 (Figure 4). The results showed that 9% of studied individuals were assigned to unique ancestry corresponding to the Iquitos, Criollo, and Contamana genetic groups from which some of the accessions included in the analysis have a recognized ancestry (e.g. IMC-67, C58, SCA-6). An additional unique group of three Colombian accessions (EBC-06, EBC-09 and EBC-58) was also discovered. The other 91% of the individuals presented a mixed ancestry.

**Figure 4.**
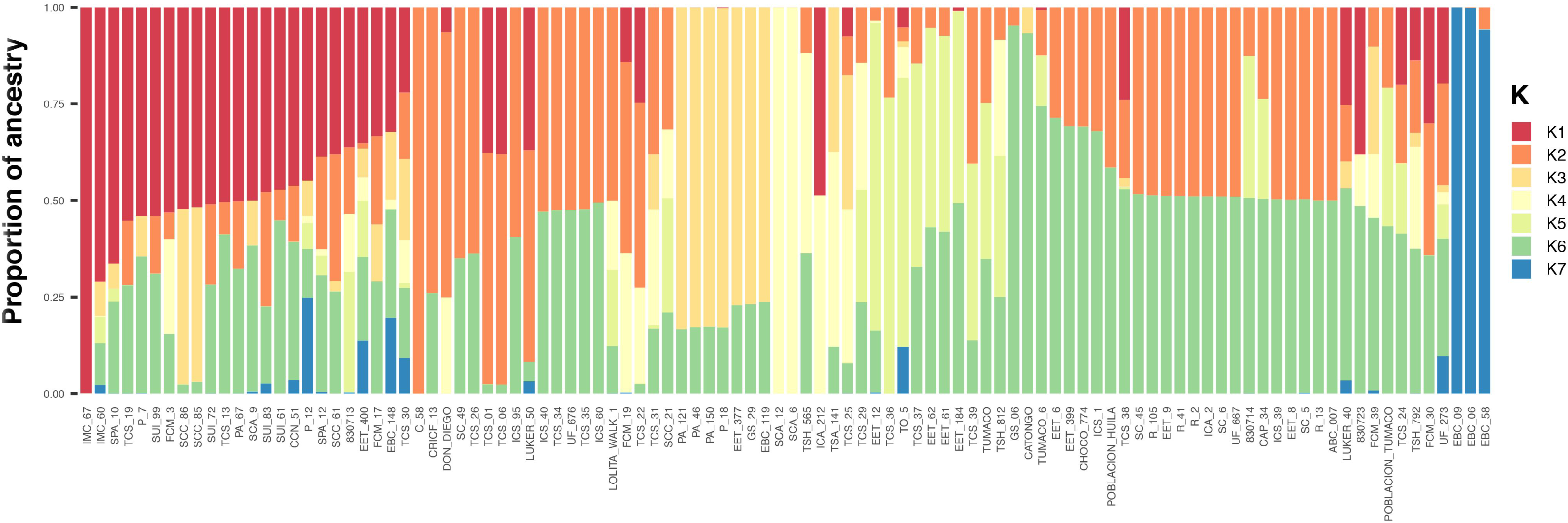
Population structure of *T. cacao* Agrosavia’s germplasm using SNPs mapped against Criollo genome. The best K was 7. The colour in each bar corresponds to the probability of a genotype to belong to an assigned group.

### Phenotypic data

During the evaluation of the 102 accessions, we observed an imbalance between the production of healthy and infected pods. In fact, the percentage of pods that reached harvest was 20.77% whereas 79.23% of pods were affected by FPRD. The pods infected by WBD was very low during all the harvest periods.

Correlations between phenotypic variables ranged from 0.12 to 0.54 and are shown in Figure S2. The highest correlation (0.54) was found between flower cushion brooms and deformed branches infected by WBD, possibly due to an indiscriminate preference of the pathogen for either of the two plant structures. The WBD variables (infected pods, flower cushion broom, and deformed branches) were also positively correlated. In contrast, symptoms of FPRD and WBD diseases are weakly correlated. Finally, as expected, disease variables did not show a strong correlation with productivity trait (healthy pods).

The ANOVA resulted in significant variation between genotypes for all the analyzed traits (observed at a level of *p ≤ 0.001*); suggesting that the accessions presented in the collection are highly diverse (Table 2).

**Table 2.**
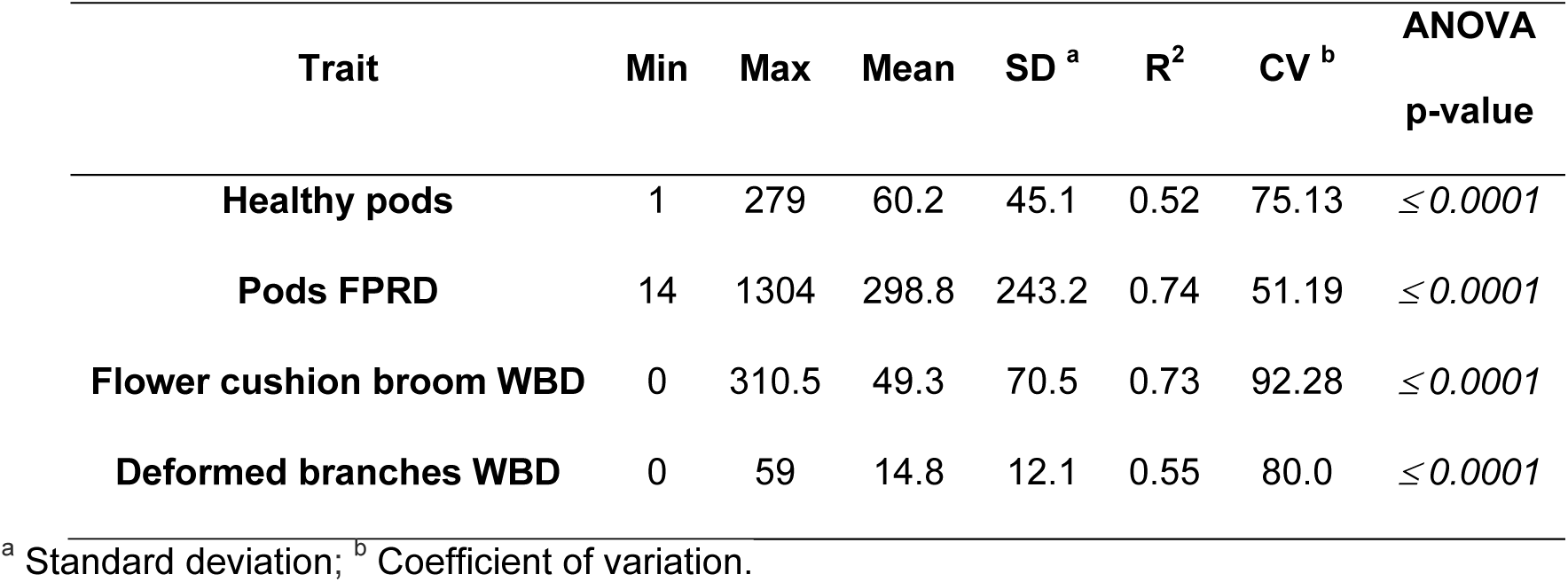
Means, standard deviation, coefficients of variation, and ANOVA’s test for the four tested variables.

| **Trait** | **Min** | **Max** | **Mean** | **SD <sup>a</sup>** | **R<sup>2</sup>** | **CV <sup>b</sup>** | **ANOVA**<br>**p-value** |
| --- | --- | --- | --- | --- | --- | --- | --- |
| **Healthy pods** | 1 | 279 | 60.2 | 45.1 | 0.52 | 75.13 | $\leq 0.0001$ |
| **Pods FPRD** | 14 | 1304 | 298.8 | 243.2 | 0.74 | 51.19 | $\leq 0.0001$ |
| **Flower cushion broom WBD** | 0 | 310.5 | 49.3 | 70.5 | 0.73 | 92.28 | $\leq 0.0001$ |
| **Deformed branches WBD** | 0 | 59 | 14.8 | 12.1 | 0.55 | 80.0 | $\leq 0.0001$ |

<sup>a</sup> Standard deviation; <sup>b</sup> Coefficient of variation.

According to the analysis of the four harvest periods, accessions GS-29, FCM-39, and EET-8 produced the higher number of healthy pods (Table S2). The genotypes CRICF-13, EBC-06, EBC-09, and SUI-72 were less affected by FPRD as shown by the AUDPC values (Table S2). The floral cushion of genotypes EET-377, SCC-85, SCC-86 and UF-273 were less affected by WBD. The genotypes EET-377 and SCC-85 also showed low number of branches affected by WBD, as well as the genotypes SUI-99, and FCM-19 (Table S2).

Based on the conglomerate analysis, the cacao accessions were divided into three main groups with different levels of productivity and susceptibility/resistance responses to WBD and FPRD (Figure 5). The first group (I) consisted of 14 accessions highly susceptible to pathogens with the highest values of infected pods by FPDR and organs infected by WBD. The second group (II) comprised 35 accessions with higher production of healthy pods and, (III) contained 53 accessions that presented the lowest values for the traits related to diseases (Table 3).

**Figure 5.**
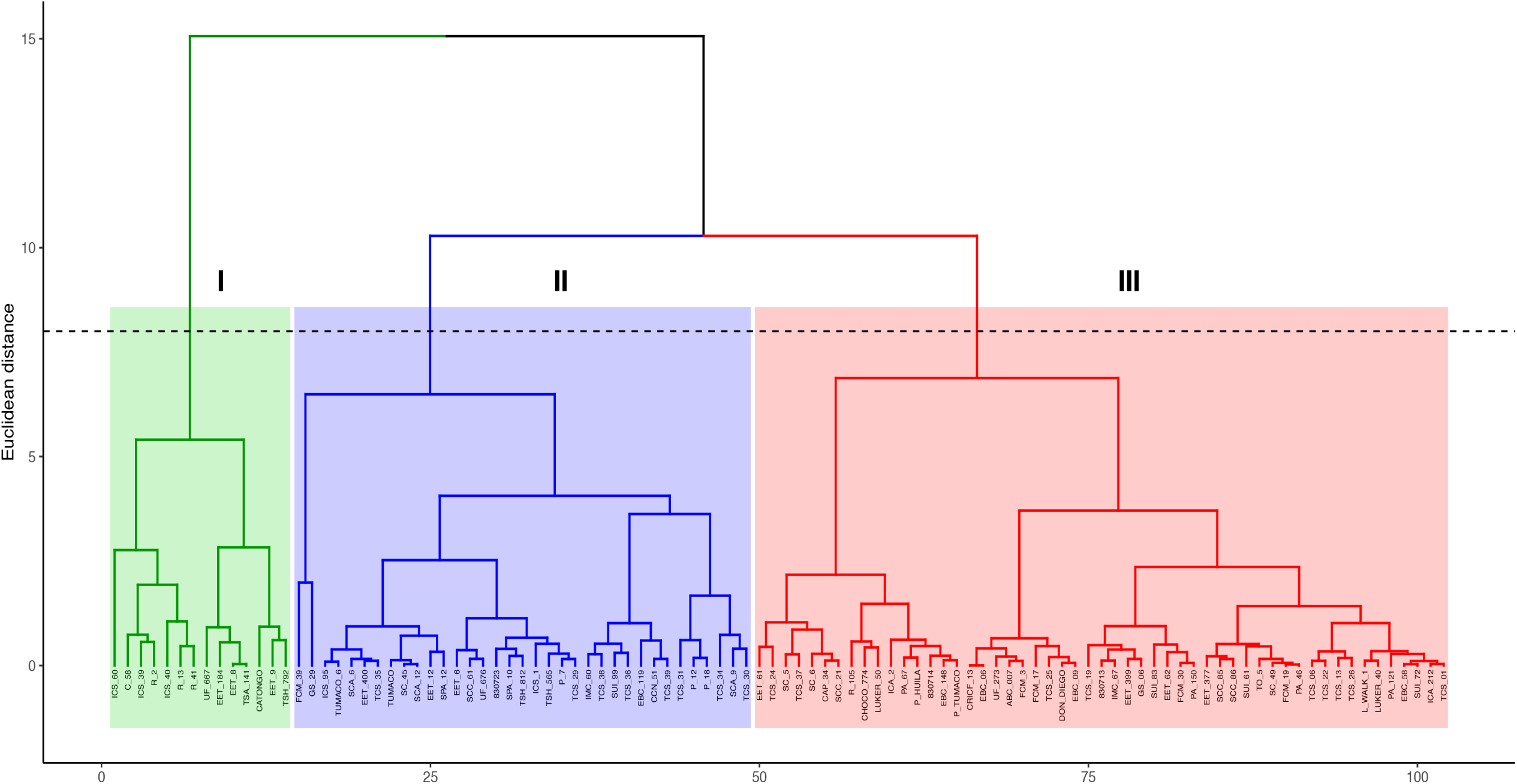
Conglomerate analysis of phenotypic data. Principal component analysis conducted using as variables the total counts of healthy pods and AUDPC variables.

**Table 3.** Statistics of evaluated traits for each group.

| **Trait** | **Cluster** | **Mean** | **SD** | **CV** | **Min** | **Max** |
| --- | --- | --- | --- | --- | --- | --- |
| **Healthy pods** | Cluster 1 | 79.86 | 38.46 | 48.16 | 33 | 165 |
|  | Cluster 2 | 87.97 | 53.48 | 60.79 | 12 | 279 |
|  | Cluster 3 | 36.68 | 22.73 | 61.96 | 1 | 95 |
| **Pods with**<br>**FPRD** | Cluster 1 | 523.21 | 263.84 | 50.43 | 264.5 | 1304 |
|  | Cluster 2 | 422 | 253.75 | 60.13 | 116 | 1225.5 |
|  | Cluster 3 | 158.15 | 107.08 | 67.71 | 14 | 414.5 |
| **Flower with**<br>**cushion WBD** | Cluster 1 | 187.61 | 91.77 | 48.92 | 65 | 310.5 |
|  | Cluster 2 | 35.29 | 27.73 | 78.59 | 1 | 97 |
|  | Cluster 3 | 22.09 | 31.84 | 144.11 | 0 | 142 |
| **Branches with**<br>**WBD** | Cluster 1 | 32.82 | 14.39 | 43.85 | 8.5 | 59 |
|  | Cluster 2 | 13.33 | 7.05 | 52.93 | 1 | 37.5 |
|  | Cluster 3 | 11.09 | 9.96 | 89.78 | 0 | 41.5 |

### Association analysis

The SNPs with an association to the analyzed traits were distributed in 5 of the 10 cacao chromosomes (Table 4). The Q-Q plots supported the association of SNP with the traits significantly (*p ≤ 0.005*) and suggested that population structure was adequately controlled in the GWAS model (Figure S3). All the associations were significant (*p ≤ 0.05*) before the FDR correction; however only the associations for flower cushions with WBD were maintained after the correction. The non-significant p-values using FDR correction is probably due to the reduced sample size of the collection.

**Table 4.**
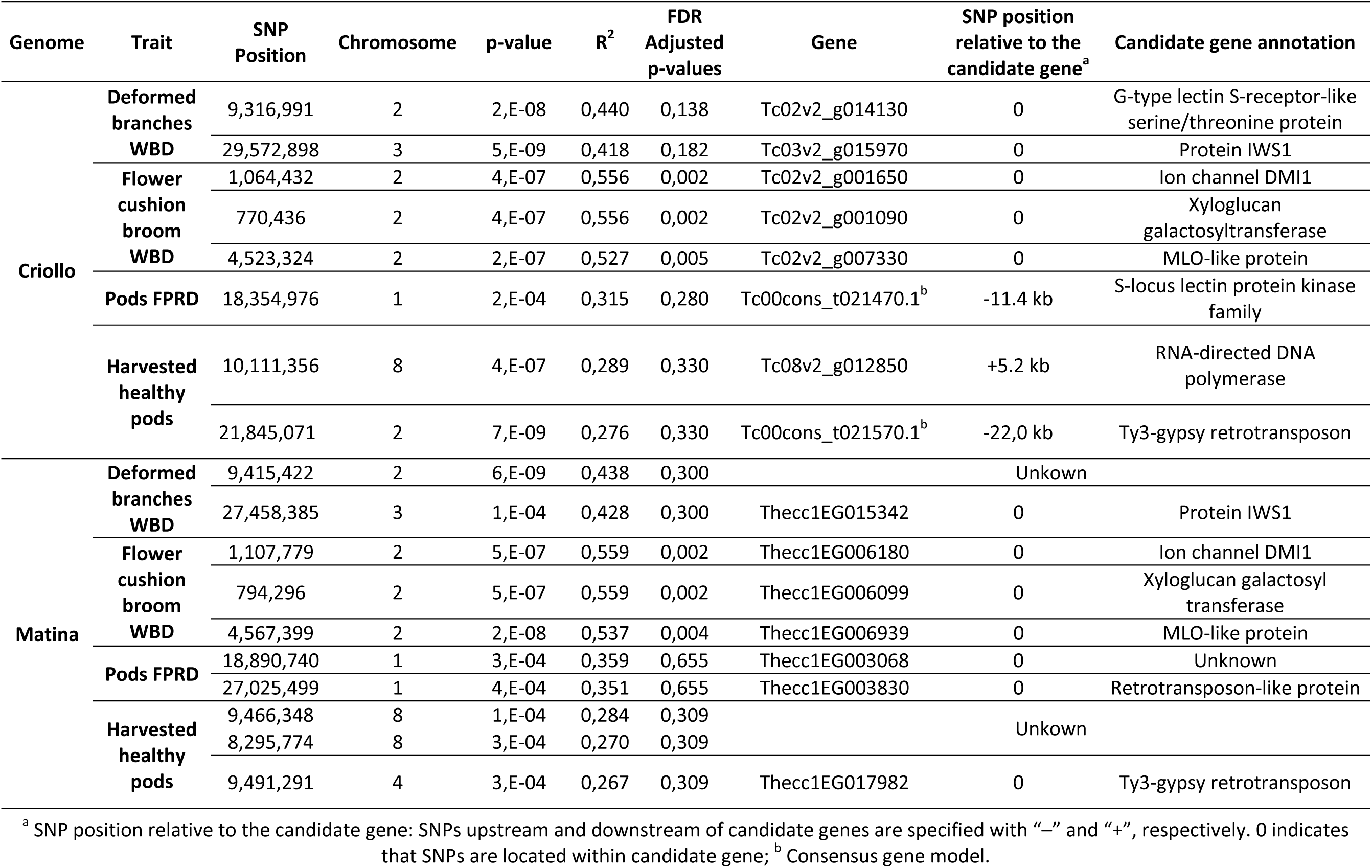
Significant marker–trait associations for evaluated traits.

Manhattan plots showing the log_10_(*p*)-values for the markers according to their positions in each reference chromosome are shown in Figure 6. The GWAS identified four loci on chromosome 2 and one on chromosome 3 associated to WBD. The annotation of these loci was exactly the same using both genomes as reference (Table 4).

**Figure 6.**
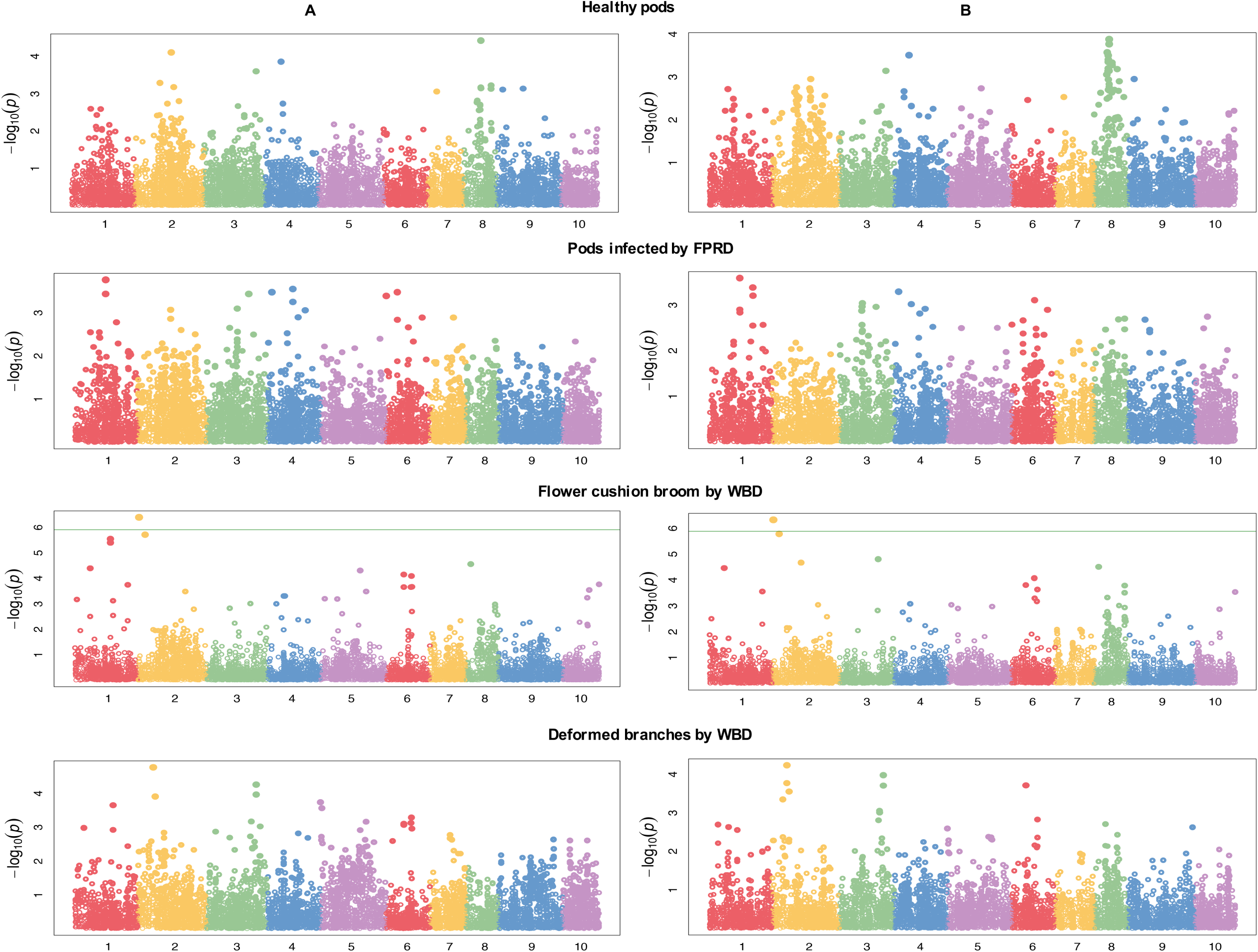
Manhattan plots of marker-trait associations for productivity, WBD and FPRD for both reference genomes. **A.** Criollo. **B.** Matina.

**Figure 7.**
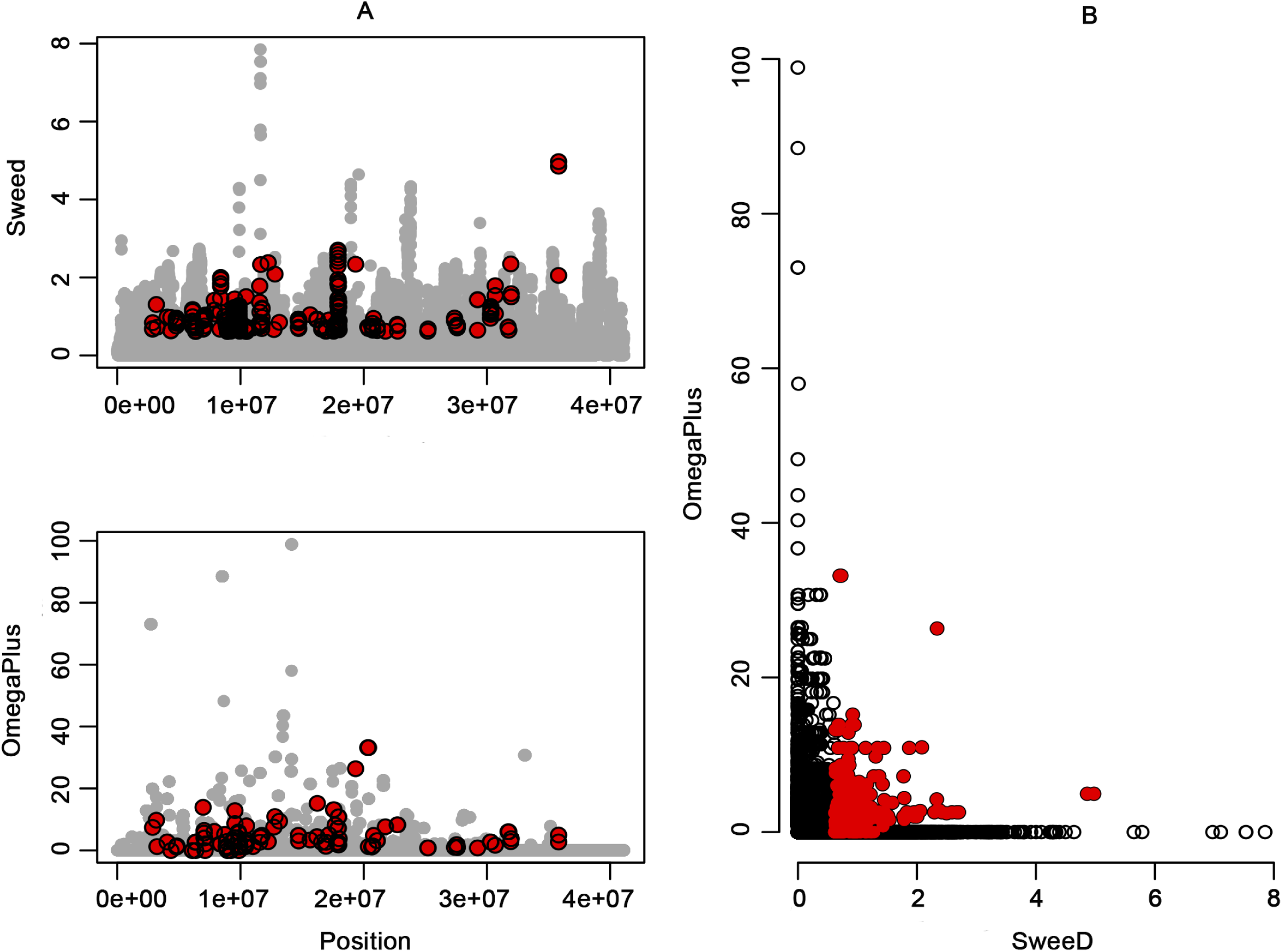
Selective sweep analysis for each chromosome. **A**. The x axis denotes the position on the chromosome, and the y axis shows the CLR evaluated by SweeD (upper panel) and the ω statistic (bottom panel) evaluated by OmegaPlus. **B.** The joint plot for SweeD and OmegaPlus. Red dots denote outliers at a significance level of 5%.

Loci related to FPRD were located on chromosome 1 using both genomes as reference. The SNP S1_18354976 was upstream 11.4 kb to the transcript Tc00cons_t021470.1 of the gene consensus model of the Criollo genome and probably correspond to the same SNP, S1_27025499, identified with the Matina genome. A second SNP located in the gene Thecc1EG003830 was identified using Matina (Table 4).

For productivity, three genomic regions were identified on chromosomes 2, 4, and 8. Two candidate genes found on Criollo genome were not located in the coding region, but they were located downstream of the gene Tc08v2_g012850 and upstream of the Tc00cons_t021570.1 transcript. Based on the Matina genome, two associated SNPs were located in chromosome 8 in genes without a functional annotation and one SNP was located on chromosome 4 (Table 4).

The heatmaps presented in figure S4 showed LD measures for chromosomes 1, 2 3, 4, and 8 which present genomic regions associated to the studied traits. As expected in an outcross species like cacao, the LD blocks were small for all chromosomes in accordance to the rapid LD decay (Figure 3).

### Identification of regions under selection

The presence of positive selection was tested by scanning the genome for i) regions of reduced variability and ii) local pattern of high linkage disequilibrium. The selective sweeps were identified by two software, SweeD and OmegaPlus; the consensus results for both software were considered the genes under positive selection. The highest number of genes was identified in chromosome 3 (39) and the lowest was identified in chromosome 1 (5) (Table S4).

The genes under selection in chromosome 1 allowed to identify a selective sweep on a region comprised between 11.1 and 22.6 million base pairs (Mbp), in which the previous associated gene Tc00cons_t021470.1 was located (Table 4). The region of chromosome 2 under selection was near to the gene Tc02v2_g014130 related to the response to WBD and related to the Tc00cons_t021570.1 transcript associated to healthy pods. The selective sweep on chromosome 3 covered a genomic region from 8.8 to 36 Mbp in which is located the gene Tc03v2_g015970 related to the response to WBD. The region in chromosome 4 under selection was not related to any of the candidate genes found by the GWAS. Finally, the selective sweep for the chromosome 8 was comprised from 9.3 to 18 Mbp in which the gene Tc08v2_g012850 associated to healthy pods was found.

## DISCUSSION

### Genotypic analysis

The main objective of the cacao breeding program in Colombia is to accumulate favorable alleles for productivity and disease resistance (Rodriguez-Medina *et al*. 2019). However, the development of an improved cacao variety could take several years because cacao is a perennial species with long juvenile stages. In order to accelerate the breeding programs, it is necessary to identify genes or molecular markers associated to genomic regions involve in disease resistance or productivity in order to select promising materials at juvenile stages. Previous studies have led to the identification of QTLs or candidate genes for productivity and for resistance to FPRD and WBD in cacao using bi-parental mapping populations with molecular markers such as SSRs, RAPDs and AFLPs (Brown *et al*. 2005; Faleiro *et al*. 2006; Queiroz *et al*. 2008) or using germplasm collections with NGS technologies such as SNP arrays (Romero-Navarro *et al*. 2017; McElroy *et al*. 2018). However, those studies present two disadvantages. First, the use of bi-parental populations produces a low mapping resolution due to the evaluation of a limited number of recombination events (Korte and Farlow 2013). The second disadvantage is to use a SNP array to analyze a germplasm collection does not allow to identify new genetic variants (Ganal *et al*. 2012).

To solve the mentioned disadvantages, we generated sequencing data using GBS, a technology developed for crop plants which is based on reducing the complexity of the genome by its fragmentation through restriction enzymes (Elshire *et al*. 2011; Ganal *et al*. 2012; Romay *et al*. 2013). This method is simple, reproducible and a high number of samples can be multiplexed, being suitable for population studies, germplasm characterization, genetic improvement and trait mapping in a variety of diverse organisms (Davey *et al*. 2011). The analysis of GBS data in a GWAS takes advantage of the historical recombination events accumulated over thousands of generations, resulting in a high-resolution mapping (Brachi *et al*. 2011; Xu *et al*. 2017).

Originally, GBS used a single digestion of enzymes of frequent cut such as *Pst*I and *Apek*I (Elshire *et al*. 2011). However, the method of single-enzyme digestion has the disadvantage of producing a high number of short fragments and low variability in the number of reads obtained per individual (Poland *et al*. 2012). To address this issue, a study conducted by Cooke *et al*. (2016) demonstrated that the use of enzymes that cut far from the recognition site called cut smart enzymes, such as *Bsa*XI, increases the diversity of the GBS library. Osorio-Guarín *et al*. (2018) proposed a modified protocol in cacao using a single enzyme digestion with *Bsa*XI to generate a great number of informative SNPs. In our study, the *in silico* digestions of the two cacao reference genomes showed that the digestion using two enzymes (*Bsa*XI and *Csp*CI) allowed to recover 3 million bp more than the study of Osorio-Guarín *et al*. (2018). Also, SNP calling recovered a higher number of SNPs markers than the datasets obtained by Osorio-Guarín *et al*. (2018) with single digestion and by Lachenaud *et al*. (2018) with double-digestion using *Pst*I and *Mse*I. The results of the present study indicate the suitability of using a double-digestion for GBS libraries for cacao using cut smart enzymes.

The genetic analysis showed that the cacao population conserved in the Agrosavia germplasm has a medium to high level of genetic diversity, that are equivalent to other studies assessing the genetic diversity of cacao (Ji *et al*. 2013; Cosme *et al*. 2016; Gopaulchan *et al*. 2019; De Wever *et al*. 2019). However, Osorio-Guarín *et al*. (2017) found a higher heterozygosity values analyzing 700 accessions of the Agrosavia’s germplasm collection and the cacao genetic groups proposed by Motamayor *et al*. (2008) were recovered in this study. Although we used the same collection, we analyzed only 229 accessions, causing lower results in heterozygosity and different population structure results.

### Phenotypic analysis

One important goal of this study was to identify accessions in the germplasm collection with high productivity or disease resistance response to FPRD and WBD. The evaluation during the four harvest periods allowed to identify three accessions, GS-29, EET-8, and FCM-39 with high productivity (Table S2). The accession GS-29 from Grenade was previously identified as a high productivity clone (Hunter 1990). The EET-8 accession from Ecuador was highlighted in a study conducted by Aranzazu *et al*. (2009) due to its high yield of 1.500 kg/ha and the FCM-39 accession is part of the germplasm bank and is a cultivar not used commercially.

As expected the incidence of FPRD was high because the germplasm collection is conserved at the research center La Suiza, located at the Magdalena Valley region, which is considered the diversity center for *Moniliopthora roreri* (Jaimes et al. 2016). Four accessions, SUI-72, CRICF-13, EBC-06 and EBC-09, showed promissory results of resistance against *M. roreri* (Table S2). The accession SUI-72 has additional qualities reported by Parra and Duarte (2007), such as a grain index of 1.73 and a pod index of 22. The accessions CRICF-13, EBC-06 and EBC-09 are Colombian native genotypes not used commercially and further studies are necessary to confirm their resistant response.

The incidence of WBD was significantly lower than FPRD which it is probably due to the fact that in Colombia during 2016 and 2017 the “El Niño” phenomenon caused adverse conditions for the disease development that requires highly humid conditions and rain all year around for its fast spread and survival (Purdy and Schmidt 1996). The evaluation of variables related to WBD allowed to identify six tolerant accessions (SCC-85, SCC-86, SUI-99, EET-377, UF-273, and FCM-19) selected by the Agrosavia’s breeding program (Table S2). The SCC genotypes (Arguello *et al*. 1999) are elite cultivars selected from regional hybrids from the department of Santander (Colombia), while accessions SUI were sampled in the department of Antioquia (Colombia) and are cacao genotypes probably introduced from Central America. The EET-377 genotype has an Ecuadorian origin and is derived from the cross between EET-156 and Scavina 6 (End et al. 1992). Scavina 6, member of the Contamana ancestral group (Motamayor *et al*. 2008), was previously identified as a resistant genotype against WBD in Brazil (Royaert *et al*. 2016). The UF-273 has been previously reported as resistant genotype to FPRD (Romero-Navarro *et al*. 2017). Finally, the FCM-19 accession is part of the germplasm bank and is a cultivar not used commercially.

### Association analysis

Biparental mapping studies from several F_1_ and F_2_ populations reported QTLs for disease resistance to FPRD on chromosomes 1, 2, 5, 7, 8, 9 and 10 (Faleiro *et al*. 2006; Brown *et al*. 2007; Queiroz *et al*. 2008; Lanaud *et al*. 2009; McElroy *et al*. 2018) and for WBD on chromosome 1, 2, 4, 6, 7, 8, and 9 (Brown *et al*. 2005; Lanaud *et al*. 2009; McElroy *et al*. 2018). Besides, Royaert *et al*. (2016) identified seven candidates genes related to WBD and Romero-Navarro *et al*. (2017) reported six genes related to FPRD. In this study, associated loci to disease resistance traits were found on the same previously reported chromosomes; however, the SNPs were located in different genomic regions.

For WBD resistance response traits, three statistically significant genes were found. The first SNP was related to G-type lectin S-receptor-like serine/threonine protein (GsSRK), related to receptor-like protein kinases (*RLKs*) gene family. In plants, *RLKs* present important roles in pathogen resistance induced after the activation of the recognition receptors of microbe-associated molecular patterns (Singh and Zimmerli 2013). Previous studies showed that *GsSRK* showed plant defense function in *Nicotiana glutinosa* (Kim *et al*. 2009) and *Capsicum annuum* (Kim *et al*. 2015), and they are also co-expressed with BIR2 kinase that confers resistance to *Arabidopsis thaliana* (Blaum *et al*. 2014). The second gene associated was the (Doesn’t Make Infections 1) *DMI1* related with ion channels (Zimmermann *et al*. 1999). Studies with ion channels in species such as tomato have demonstrated their possible relationship with plant defense (Zhang *et al*. 2018). The last identified gene was related to mildew resistance Locus O (*MLO)* protein. The *MLO* is a gene family specific to plants and plays significant roles in the resistance to powdery mildew and response to a variety of abiotic stresses, plant growth and development (Liu *et al*. 2017). Studies on *MLO* genes have demostrated that confers resistance to species such as *Arabidopsis* and tomato (Acevedo-Garcia *et al*. 2014). Other candidate genes were also found for WDB resistance (*IWS1* and *Xyloglucan galactosyltransferase*), but further studies are necessary to determine their role in plant defense.

Two candidate genes were related to resistance response against FPRD. The first one was associated to S-locus lectin protein, member of *RLKs* gene family which are related to plant immunity (Singh and Zimmerli 2013) as explained before. The second gene was related to retrotransposons and according with an study conducted by Zervudacki *et al*. (2018), certain retrotransposons behave like an immune-responsive gene during pathogen defense in *Arabidopsis*.

QTLs for productivity have been previously reported on chromosomes 1, 2, 4, 5, 9, and 10 (Clement *et al*. 2003). Recently, McElroy *et al*. (2018) reported QTLs related to fresh weight of cacao pods on chromosomes 1, 3, 7, 8, 9, and 10. Previous studies on healthy pods did not find candidate genes related to the trait (Romero-Navarro *et al*. 2017). In this study, we found two candidate genes related to the healthy pod trait (Tc08v2_g012850 and Tc00cons_t021570.1); however, further studies are necessary to confirm its role in cacao productivity.

### Selective sweep analysis

Selective sweep analysis were conducted in order to identify nucleotide variation that can be associated to beneficial traits for plant adaptation (Stephan 2010). Recent studies demonstrated that genomic regions that exhibit selection signatures are also enriched in genes associated to biologically important traits (Wen *et al*. 2015; Xie *et al*. 2015). Thus, the detection of selective sweeps and GWAS and its comparison could help to unravel the genetic structure of complex traits (Chen *et al*. 2016). In our study, some of the associated genes to studied traits were located in regions under positive selection, probably indicating that natural selection is acting in accessions occurring in regions with natural incidence of FPRD and WBD. However, further studies on evolution and domestication processes are necessary to validate these hypotheses.

Our findings expand previous genetic mapping efforts and allow to increase the mapping resolution of the regions responsible for productivity and disease resistance traits. Leveraging this information could contribute to select promising accessions in juvenile stages in greenhouse conditions, and consequently reduce the breeding cycles.

### Conclusions

This research reported the first GWAS study of disease resistance trait for WBD and FPRD in cacao based on SNPs produced by the GBS technique. In total, two candidate genes related to productivity and seven related to disease resistance response to FPRD and WBD were detected. We also identified a total of 10 promissory accessions that presented a degree of resistance response and three accessions with promising values for productivity. The present work provides new knowledge on genomic regions involved in the productivity and disease resistance response that after functional validation could be useful in MAS for cacao breeding programs.

## ACKNOWLEDGEMENTS

The authors are grateful to Marco Antonio Lopez to provide the script for the LD heatmap analysis.

